# Tracing the origin of Finnish gelsolin amyloidosis using haplotype sharing trees

**DOI:** 10.64898/2026.02.11.705340

**Authors:** Osma S. Rautila, Sari Atula, Tuuli Mustonen, Eeva-Kaisa Schmidt, Miko Valori, Roberto Colombo, Juha Kere, Karri Kaivola, Pentti J. Tienari

## Abstract

Finnish gelsolin amyloidosis (AGel amyloidosis) is an autosomal dominant systemic amyloidosis caused by GSN c.640G>A p.D187N (rs121909715) founder variant. The disease was first described in 1969, and it was hypothesized that the Finnish patients share a common ancestor dating back to the 14th century. The link between two Finnish regions with high AGel incidence (Kanta-Häme and Kymenlaakso) has been hypothesized to have occurred in 1365 by a settler moving from Kanta-Häme to Kymenlaakso. Here, we used haplotype sharing tree (HST) to analyze Finnish AGel amyloidosis haplotypes to trace the geographic origin of the variant. We also estimated the time from the most recent common ancestor (MRCA) using single nucleotide polymorphism and short tandem repeat data. The HST -based analyses leveraging AGel amyloidosis cohorts from different Finnish geographic regions indicated, that the variant more likely appeared first in Kymenlaakso, not Kanta-Häme, contrary to the original hypothesis. The MRCA estimates for Finnish AGel ranged from 15 to 40 generations using four different methods, the mean of all estimates (27 generations) dated back to the 14th century. Thus, the data supports the original hypothesis on the variant’s spreading temporally, but not geographically. These results illustrate the use of HSTs in the analysis of haplotype structures and in tracing the ancestry of a founder variant.

## INTRODUCTION

Finnish gelsolin amyloidosis (AGel) [MIM: 105120] is an autosomal dominant systemic amyloidosis. The first symptom is usually corneal dystrophy, often followed by a slowly progressing facial nerve paresis and cutis laxa.^**1,2**^ AGel is caused by *GSN* [MIM: 137350] c.640G>A p.D187N (chr9:121310819, GRCh38) variant,^**3,4**^ which was initially discovered in the polypeptide,^**5,6**^ and then demonstrated in DNA sequence analysis.^**6,7**^ The variant in Finland has a founder haplotype,^**8**^ and the variants in Finland and e.g. Japan are known to have an independent origin.^**9,10**^ There is also another variant G>T at the same position,^**1,11,12**^ and in recent years, many more *GSN* variants have been discovered.^**13–20**^

AGel was first described in Finland in 1969,^**21**^ and the highest incidences have been reported in two Finnish regions: Kanta-Häme and Kymenlaakso.^**22**^ Based on population registers, it was hypothesized the patients shared a common ancestor dating to the 14th century when Finland was a part of the Kingdom of Sweden. Coincidentally, during a visit to Finland on the 16th of February in 1365, the King of Sweden, Albert of Mecklenburg, had granted a letter of protection to a settler Matti Orava who had moved his community from Kanta-Häme to Kymenlaakso, but had faced aggression from local peasants.^**23**^ Under the protection of the King, Orava founded the village of Oravala in Kymenlaakso. One hypothesis was, that this founding could be the potential link between patients from Kanta-Häme and Kymenlaakso.^**24**^ If this hypothesis were true, the variant would then have originated first in Kanta-Häme prior to 1365 and later moved to Kymenlaakso.

Here, we set out to explore whether genetic evidence supports this hypothesis. Haplotype sharing trees (HSTs) constructed from phased SNP array data for the study of identity-by-descent (IBD) segments was previously shown to be a good fit in the Finnish population.^**25**^ Here, we expand the use of HSTs to tracing the origin of the *GSN* c.640G>A variant in Finland based on haplotype similarities.

## METHODS

### Cohorts, quality control and phasing

Our cohort contained 62 nuclear GSN families from our previous study.^**8**^ We genotyped 78 individuals using the Illumina GSAv3 SNP array. The data was quality controlled (Supplemental Methods 1) and markers that were not on the Finnish SISU v3 reference panel were excluded.

Phasing was done using Beagle 5.3,^**26,27**^ after which 66 samples (IBD cut-off 0.1875) and 350,093 SNPs remained across all chromosomes for the MRCA analysis. For the geographical origin analysis, we included the Finnish regions which had 4 or more samples available, which resulted in a total of 53 samples.

The geographical origin for each sample was based on self-reported knowledge of ancestry.

For short tandem repeat (STR) genotyping, we selected nine AGel parent-offspring pairs, 20 unrelated AGel patients and 56 healthy controls. All AGel patients were confirmed heterozygous for *GSN* c.640G>A with Sanger sequencing. The controls did not have the variant.

Because the *GSN* variant c.640G>A is not present on the genotyping array, we selected the haplotype that shared more sequence with the rest of samples, and discarding the shorter one.^**25**^ The success of the longest haplotype -based selection was evaluated using a haplotype comparison graph (Figure S3).

This study was approved by the Institutional Review Boards of the Helsinki University Hospital (Drno 200/13/03/01/2013).

### Haplotype sharing trees

Haplotype sharing trees store all shared haplotypes starting from a predefined point in the genome into a tree-like data structure. HSTs are always majority-ordered, meaning the haplotypes in the majority of samples are stored to the left node. By following the left side of the tree (i.e. *the majority-branch*), the last shared haplotype at the end of the branch is what we call the *majority-based ancestral haplotype* (as it is dictated by the majority) (for construction details see Figure S1 and S2).

Here, we use HSTs to analyze the similarities between *GSN* haplotypes from different regions in Finland. For the analysis of *GSN* c.640G>A, using HAPTK, we construct two HSTs. One moving towards the left from *GSN*, and another moving towards the right. This allows us to analyze haplotype sharing and recombination patterns on both sides of the variant.

Under the assumption that migrations of the rare GSN variant have not happened back and forth, the haplotypes of all non-originating regions are then sampled from the region of origin. Thus, in theory, the region of origin should have the oldest and most diverse set of haplotypes.

### MRCA estimation

For MRCA estimation on the SNP array data, we used the Gamma method by Gandolfo et al^**28**^ implemented in HAPTK.^**25**^ As the starting marker for the analysis, we selected the variant nearest to *GSN* c.640G>A located at chr9:121312513 (GRCh38). For STR data -based MRCA estimation, we selected four *GSN* flanking STRs: downstream D9S195 and D9S258, and upstream D9S282 and D9S778 (Table S8). The alleles were genotyped using PCR and capillary electrophoresis (Supplemental Methods 2, Table S6 and S7). We then used three methods for MRCA estimation. First, Disease Mapping Using Linkage Disequilibrium 2.3 (DMLE+ 2.3), which implements a Bayesian LD mapping method.^**29**^ Second, Moment of Methods estimator^**30**^ for each marker using two different population growth rates 0.08 and 0.14 with the linkage disequilibrium calculated from allelic excess according to Bengtsson and Thomson.^**30**^ Third, Luria-Delbruck based model with the same parameters.^**31**^

## RESULTS

### Haplotype Sharing Trees (HSTs) and the ancestral haplotype

We first determined the core haplotype, shared by all *GSN* samples, which was 64-markers long (2,2 Mb) ranging from 119,632,253 to 121,864,249 (GRCh38). The marker alleles and their positions are shown Table S1.

The majority-based ancestral haplotype (n=66 samples) ranged from 107,626,124 to 130,233,489 (22,6 Mb) (marker alleles shown in Table S2). After deriving the majority-based ancestral haplotype, phasing was inspected by comparing all haplotypes against it. The phasing separated the variant haplotypes reliably from other (wild-type) haplotypes with minor potential switch-errors in the data (Figure S2, variant haplotypes in white, wild-type haplotypes in blue, potential switch-errors shown in arrows).

Because the c.640G>A variant is known to have separate origins in different populations,^**9,10**^ we made the HSTs with haplotypes shared by at least ten samples publicly available for future comparisons against samples of different origin.^**32**^

### Geographical analysis

The patients’ regions of birth are indicated in Figure 1A. We constructed HSTs for the 53 *GSN* haplotypes originating from these five regions. The haplotypes from Pirkanmaa (light blue) shared relatively short segments with the other regions, whereas haplotypes from the four other regions extended all the way to the end of the majority-branch (Figure 1B and C, at 3 o’clock). Physical distances shared with the majority-based ancestral haplotype are shown in Figure 1E.

**Figure 1.**
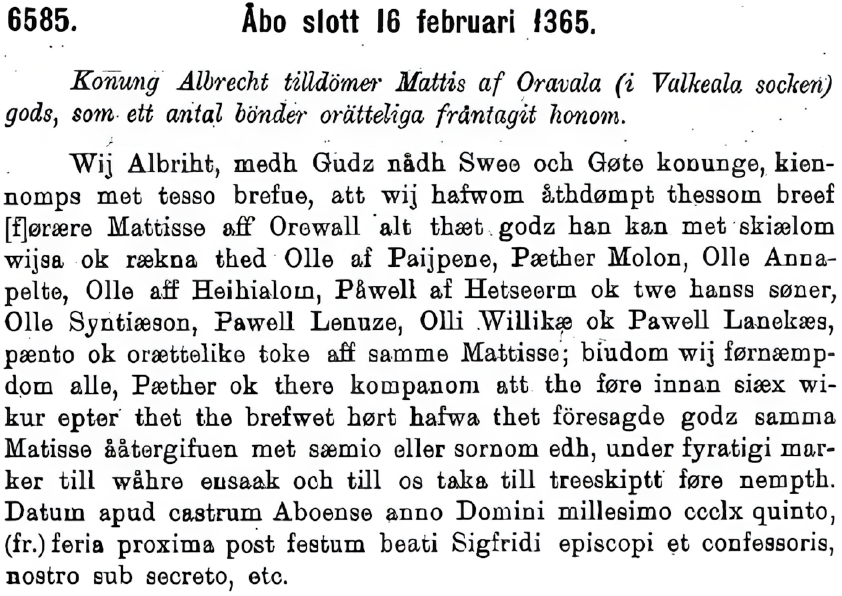
A letter from King Albert of Mecklenburg (16th of February 1365) *King Albert grants Matti of Oravala (from the county of Valkeala) the estate that a number of peasants have unjustly taken from him*. From the collection of the Finnish state archive (Finlands medeltidsurkunder VIII, p. 359).

All regions formed some level of clustering, and the region of Kymenlaakso (purple) formed the largest, and earliest branching, subtrees in both left- and right-sided HSTs (Figs 1B and 1C). To further investigate the origin of the variant, we selected the regions present near the end of the majority-branch and thus excluded Pirkanmaa which shared only short segments with the ancestral haplotype (Figs 1B and 1C). We then constructed HSTs for the regions individually and derived the regional majority-based ancestral haplotypes. The ancestral haplotypes of each region were aligned, and the consensus identities were constructed using CIAlign^**33**^ (Figure 2E). The ancestral haplotype from Kymenlaakso shared the longest consensus identities both on the left and right side (Fig 2B-D) in relation to the other regional ancestral haplotypes. To the left, the Kymenlaakso ancestral haplotype formed a majority with Karjala, and to the right with Kanta-Häme (Figure 2E).

**Figure 2.**
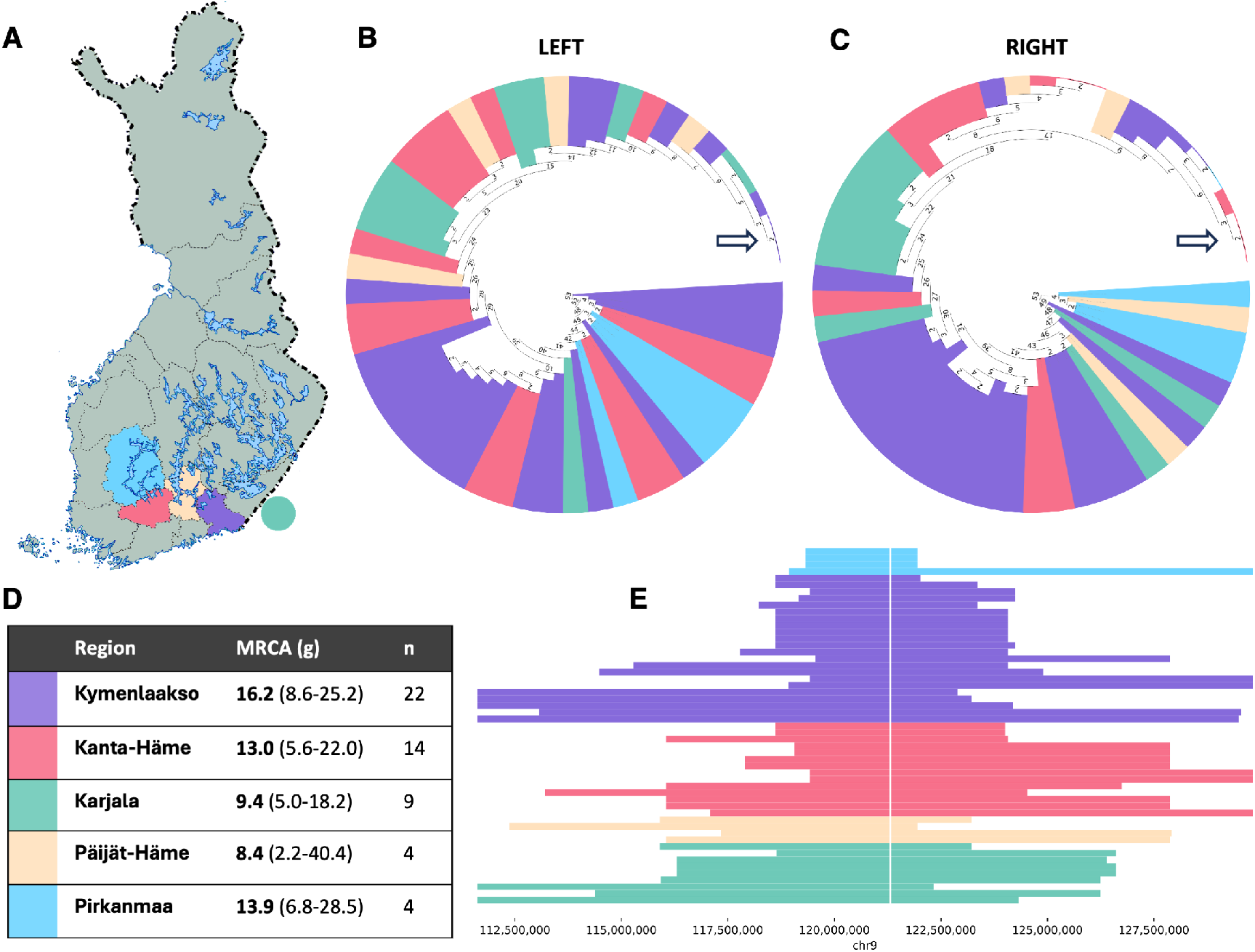
Haplotype sharing and regional MRCAs. **(A)** Map of Finland with the five regions containing over four GSN samples. The green dot represents Karjala, which was ceded to the Soviet Union after World War II. The original hypothesis proposed that Mr. Orava settled from Kanta-Häme (in red) to Kymenlaakso (in purple) around 1365 during which the GSN variant arrived to Kymenlaakso. **(B)** Left-side HST where each leaf-node is colored by the haplotype carrier’s family region. Following the left-most nodes leads to the majority-based ancestral haplotype (arrow). **(C)** As B., however right-side HST. **(D)** Table containing the names and color codes of the regions. MRCAs were estimated with the Gamma method. **(E)** Segments of the majority-based ancestral haplotype shared by each sample. Each horizontal line represents a GSN haplotype (n=53) and the color the region of origin. The GSN locus is marked by the white line. The haplotypes are ordered by length, and the groups are ordered by average length of ancestral haplotype sharing (highest at the bottom). The x-axis is windowed for visualization purposes.

Last, we analyzed how long segments individual haplotypes from each region shared in common with the regional ancestral haplotypes. Out of the larger regions, Karjala and Kanta-Häme shared the largest segments of haplotype with their own regional ancestral haplotype. Strikingly, haplotypes from all regions shared approximately equally long sequences with the Kymenlaakso ancestral haplotype. Also, haplotypes from Kymenlaakso shared 1 to 2 Mb less sequence with other regional haplotypes, indicating larger diversity within the Kymenlaakso haplotypes.

### Most recent common ancestor (MRCA) estimation of *GSN H*c.640G>A

To estimate the number of generations until the MRCA, we first used the correlated genealogy option of the Gamma method on the SNP data which resulted in an estimate of 15.2 (CI 95%, 7.8-30.1) generations. We then estimated the MRCA for individual regions (Figure 1C), which resulted in Kymenlaakso having the oldest MRCA estimate at 16.2 g (CI 95%, 8.6-25.2).

The region of Kymenlaakso also contained the largest number of samples, which could affect the estimate. To minimize this bias, we randomly downsampled the Kymenlaakso cohort to 14 samples (same size as Kanta-Häme) and performed 1k iterations of MRCA estimation leading to an average estimate of 14.5 g, which was still the highest of all the regions.

Next, we estimated the MRCA using STR data and the DMLE+ 2.3 software assuming a mean population growth rate of 0.11.

The distribution of the posterior probability of the variant age was obtained after a burn-in of 100,000 iterations followed by a run of 1,000,000 iterations. The maximum likelihood estimate corresponded to 30.9 generations (95 % credible set: 16.6–61.8 generations) (Figure 3C, Table S3). Third, using the STR data and Luria-Delbruck -based method, the MRCA estimate was 19 generations (CI 95%, 12-26). Fourth, using the STR data and Moment of Methods estimator the MRCA estimate was 42 generations (Table S4).

**Figure 3.**
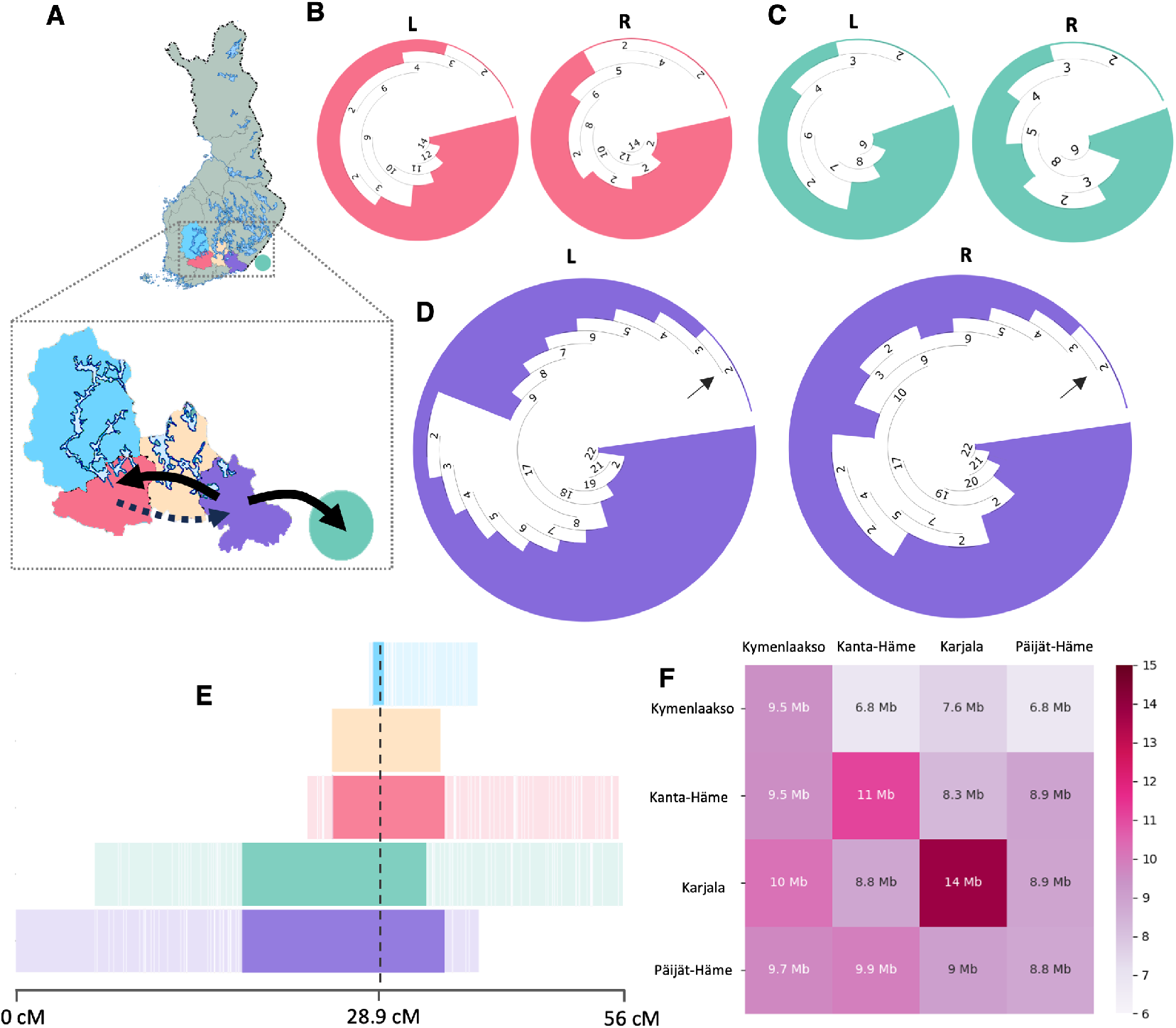
Geographical haplotype similarity. **(A)** Map of Finland showing the originally hypothesized geneflow in dotted arrow and the more likely flow in solid arrow. **(B to D)** HSTs of **(B)** Kanta-Häme (red), **(C)** Karjala (green) and **(D)** Kymenlaakso (purple). In the Kymenlaakso HST the arrows remind of the node containing the majority-based ancestral haplotype. The HST of the left side splits into two subtrees of 9 vs 8, narrowly deciding the majority. **(E)** Majority-based ancestral haplotypes as consensus identities from CIAlign. The shared consensus segments for each haplotype are shown around the GSN locus (dashed line). The lower opacity regions are the remainders of the ancestral haplotypes, and most likely represent recombined sub-founder haplotypes. The white lines indicate mismatching SNPs. The haplotypes are ordered by total consensus length around GSN with Kymenlaakso (purple) having the longest consensus. **(F)** On the top are majority-based ancestral haplotypes of each region. All the individual haplotypes from each region are compared against these haplotypes, and the average length of the shared segment is calculated. All regions share 9.5-10 Mb sequences in common with Kymenlaakso, however, haplotypes from Kymenlaakso share shorter sequences in common with the majority-based ancestral haplotypes of other regions.

Figure 4B provides the summary of the MRCA estimates and their confidence intervals. Despite differing point estimates, the confidence intervals overlapped for each estimate. The average of all estimations was 27 generations.

**Figure 4.**
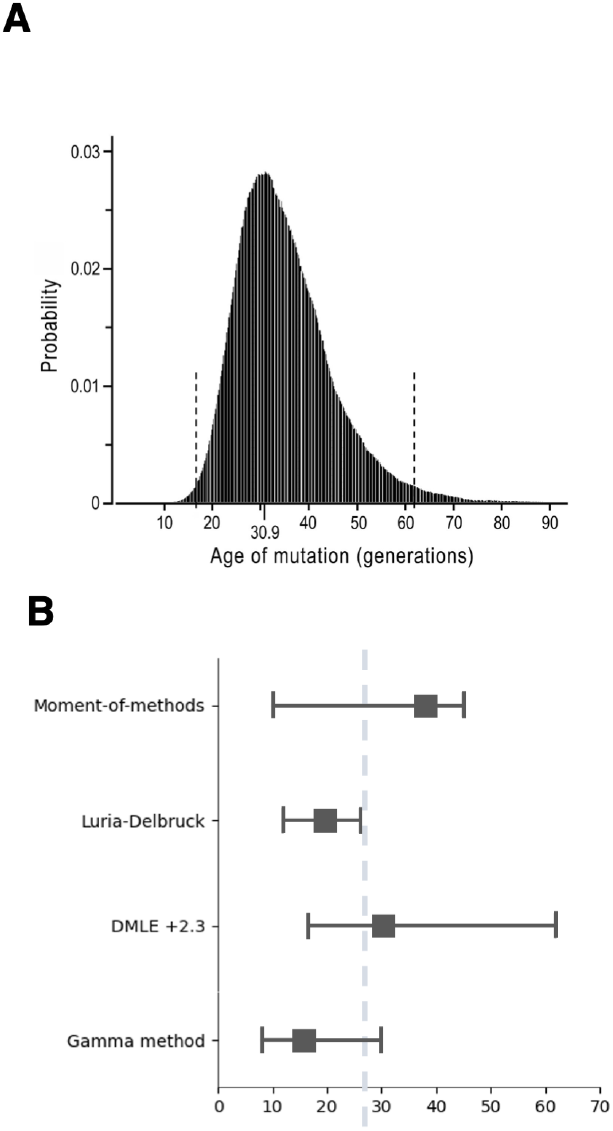
MRCA estimates. **(A)** DMLE 2.3+ Bayesian estimation of the variant age **(B)** MRCA estimates and confidence intervals per each technique. The dashed line denotes the average of all point estimates.

## DISCUSSION

Using genetic evidence, we have tackled the hypothesized link between two events. The appearance of the *GSN* c.640G>A variant (with the highest incidence in two regions: Kymenlaakso and Kanta-Häme), which based on church records could have occured in the 14th century, coinciding with the King of Sweden, Albert Mecklenburg granting estate rights in Kymenlaakso to a settler from Kanta-Häme, Matti Orava, in 1365.

Contrary to the prior hypothesis that the variant migrated from Kanta-Häme (or Karjala) to Kymenlaakso, our results provide indication that it in fact originates from Kymenlaakso. First, the regional ancestral haplotype of Kymenlaakso formed the largest consensus between the regional ancestral haplotypes. Second, haplotypes from all regions shared long segments with Kymenlaakso ancestral haplotype, but not vice versa, indicating higher diversity within the Kymenlaakso haplotypes. High diversity was also indicated by earlier branching in HSTs and the presence of Kymenlaakso haplotypes in most large branches. Third, the regional MRCA was marginally higher for Kymenlaakso. Fourth, geographically the simplest scenario is that the variant has moved from Kymenlaakso both to the west and to the east.

Thus, these actions by Mr. Orava and king Albert of Mecklenburg most likely did not contribute to the distribution of the *GSN* c.640G>A variant in Finland.

However, our results are temporally in accordance with the original hypothesis and the evidence from church records. The MRCA of the whole GSN cohort was estimated using four different approaches. The Gamma method gave the smallest estimate at 15.17 generations and the moment of methods the highest at around 42 generations. The estimates ranged from 15 to 42 generations (375 to 1050 y) and thus the MRCA could have lived around 970 - 1620 AD. The average of all point estimates was 27 generations (25 years per g, 670 y) which would correspond to year 1354.

This study has limitations. First, not all GSN families from Finland were included in this study, which may underestimate the number of generations in MRCA estimation and affect the majority-based haplotypes. Second, the MRCA analyses have wide confidence intervals, and the results should be interpreted as non-exact. Also, the Gamma method assumes a Haldane model of recombination and the phased haplotypes are susceptible to switch and genotyping errors, which would cause bias towards shorter shared haplotypes and higher MRCA estimates. Third, in the STR based methods, the number of patients was lower (n=38) and one of the five STRs was quite monomorphic indicating it was very close to the core haplotype.

Nonetheless, we demonstrate the applicability of HSTs in a cohort specific IBD-based analysis of variant ancestry.

## CODE AND DATA AVAILABILITY

HAPTK and all the scripts used in this article are available at https://github.com/xosxos/haptk. The Finnish GSN HSTs are available in Zenodo under DOI: https://doi.org/10.5281/zenodo. 14589217

## ACKNOWLEDGEMENTS

The data was phased with the THL Biobank’s SISU v3 Imputation reference panel obtained from THL Biobank. We thank all study participants for their generous participation in the FIN-RISK, Health 2000, and Migraine Family studies. We also thank the Sequencing Informatics Team, FIMM Human Genomics, University of Helsinki for the work done on the reference panel. Finally, thank you to the Finnish IT Center for Science (CSC) for providing computing resources used in this study.

This study was funded by the Finnish Medical Foundation, Finnish Cultural Foundation (Kymenlaakso), Päivikki and Sakari Sohlberg Foundation, Paulo Foundation, Sigrid Juselius Foundation, Finnish Brain Foundation, Biomedicum Helsinki Foundation, Helsinki University Hospital grants and Academy of Finland (318868).

## SUPPLEMENTAL METHODS

### 1. SNP data quality control

Genotyping data quality control was performed separately on samples processed with different arrays. First, we removed duplicated samples and related samples (proportion IBD > 0.1875), samples with discordant sex information, outlying heterozygosity rate (>3SD) and > 5% missing genotype rate. In per-variant quality-control, we included variants that had a genotyping rate of > 95%, variants in HWE (p>0.000001) and minor allele frequency >= 0.01. To harmonize genotyping array data, we included only biallelic non-palindromic SNPs and set allele coding to match the reference genome so that A1 allele was always the alternative allele regardless of minor allele frequency. After the SNP array data quality control, we excluded markers with discrepant AF to the SISU v3 reference panel or not present in the reference panel.

Then, we excluded markers with an allele count of under 3 and performed phasing using Beagle 5.3 with ne set to 20 0000 and the number of iterations set to 128.

### 2. Short tandem repeat PCR and capillary electrophoresis procedure

We selected two STRs flanking the GSN c.640G variant both down and upstream. Downstream STRs D9S195 (AFM193yg5) and D9S258 (AFM185xe3) were located at suitable distances. Upstream we selected STRs D9S282 (AFM308vb1) and D9S778 (UT7968). Next, we determined suitable primers for PCR by performing a look-up of the STRs in the BLAST database (Table S5). The DNA was run through a polymerase chain reaction and the size of the resulting DNA fragments were determined through capillary electrophoresis using standard practices. Last, the alleles were determined for each sample with GeneMapper 6.0. The PCR mix of STRs samples was done with 4.6 uL of Betaine, 2.3 uL of DreamTaq buffer, 0.4 uL of DreamTaq polymerase, 1.15 uL of reverse and forward primers, 0.7 uL dNTP and 50 ng / uL of DNA resulting to a total volume of 20 uL. The PCR reaction was performed with initial denaturation of 95 °C for 5 minutes following 29 cycles of denaturation at 95 °C for 20 s, annealing at 58 °C for 20 s and extension at 72 °C for 20 s and finalized by extension at 72 °C for 5 minutes. Depending on the intensity detected on gel electrophoresis the resulting samples were then diluted by 1:100 or 1:200 before capillary electrophoresis.

## SUPPLEMENTAL FIGURES

**Figure 5.**
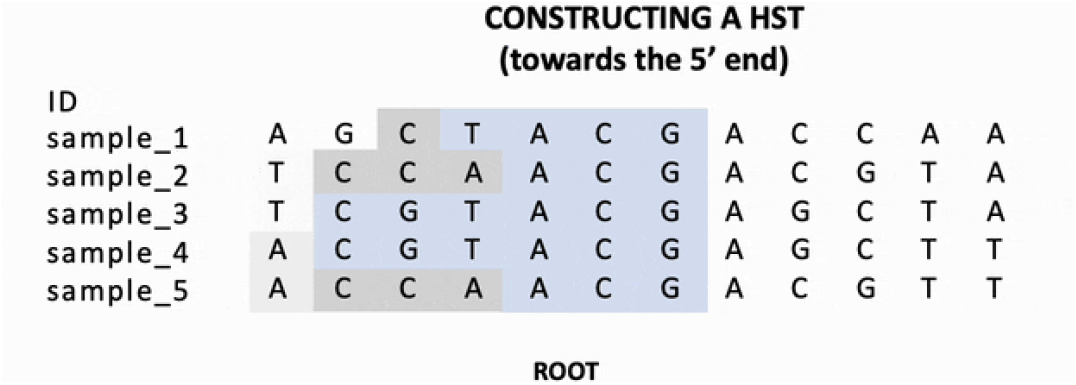
Figure S1. Haplotype sharing trees and the majority-based ancestral haplotype. Link: https://github.com/xosxos/haptk/blob/main/assets/hst.gif

**Figure 6.**
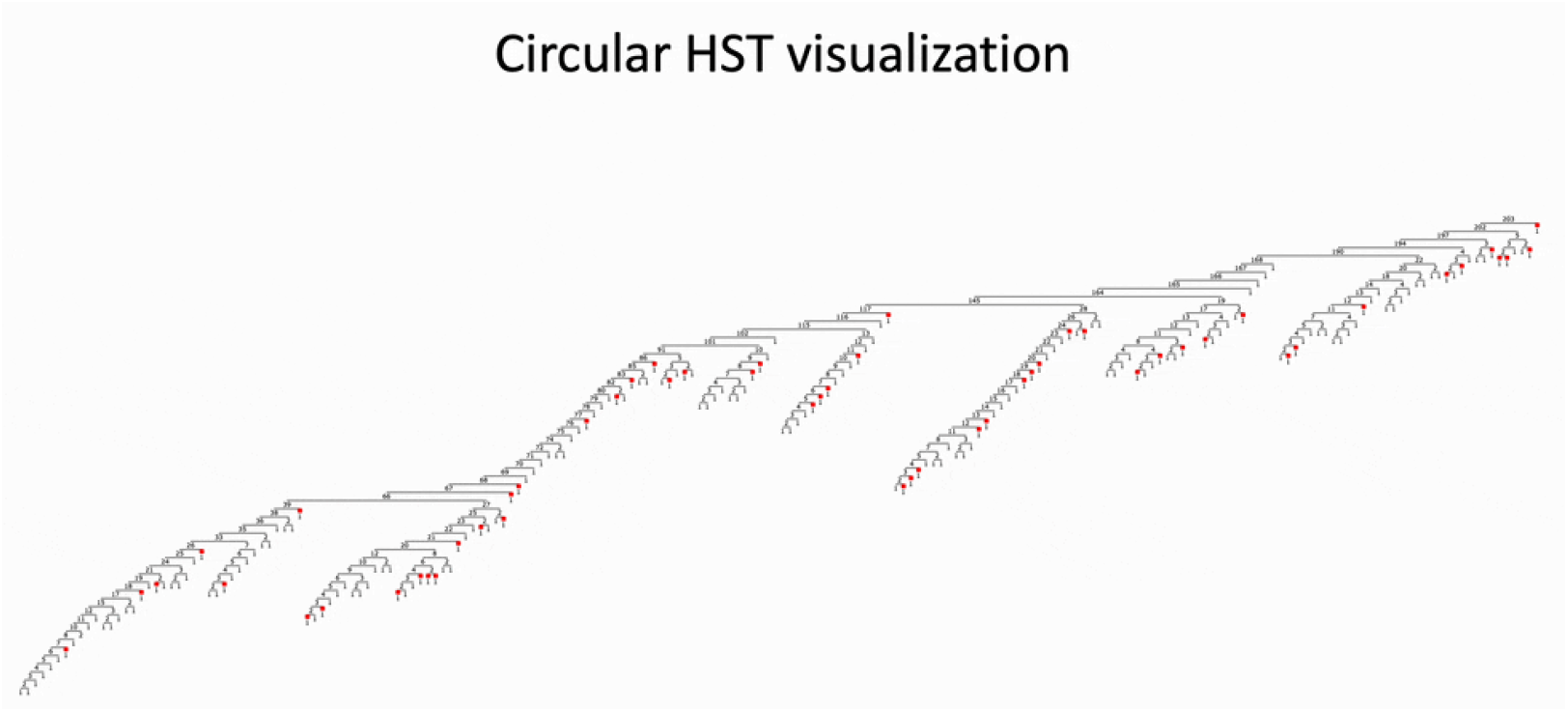
Figure S2. Constructing the circular HST representation. Link: https://github.com/xosxos/haptk/blob/main/assets/circular_hst.gif

**Figure 7.**
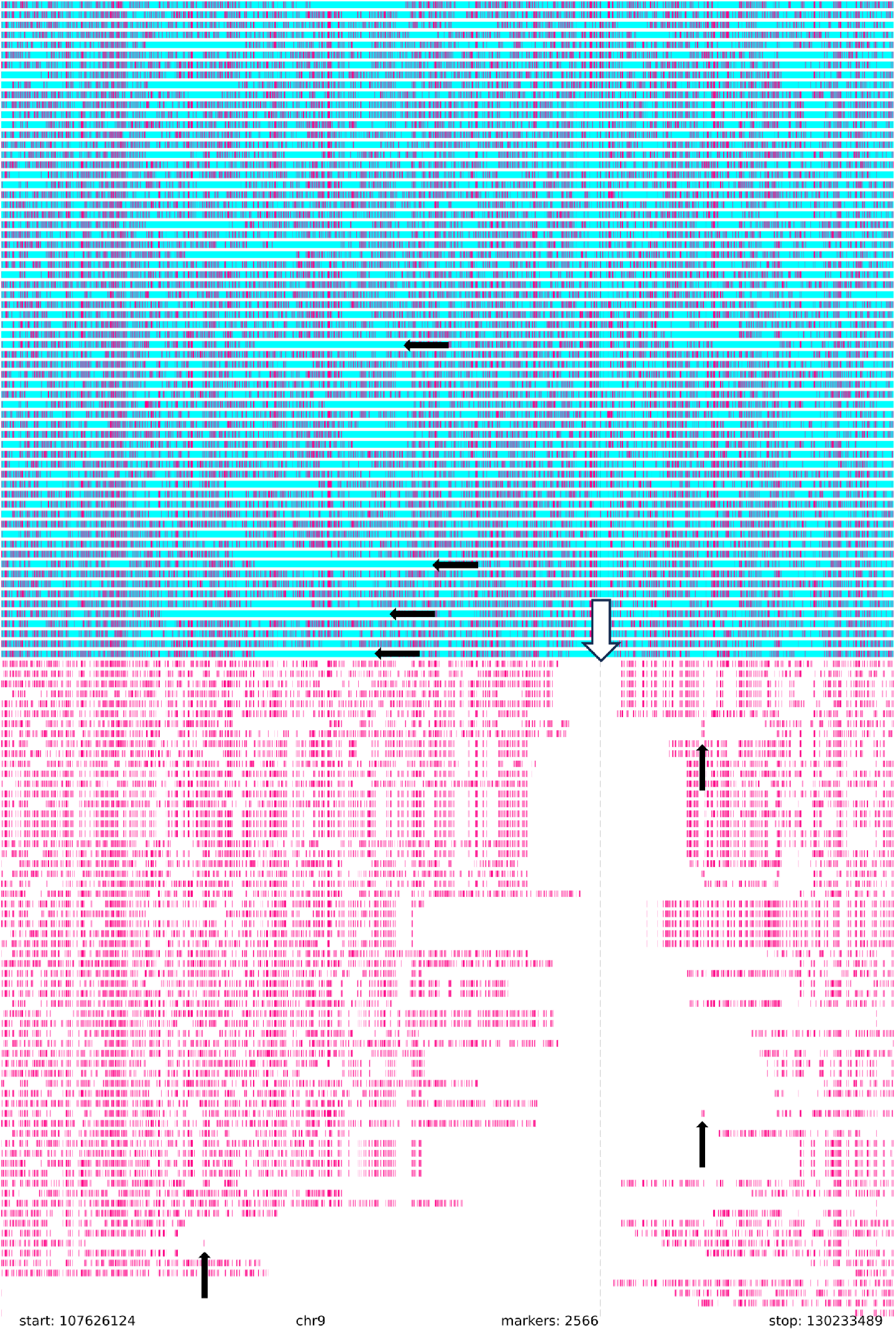
Figure S3. Haplotype selection, switch and genotyping error evaluation based on the ancestral haplotype comparison graph. The white down-arrow on the top denotes the GSN locus. The black up-arrows denote suspected genotyping errors or small switch-errors. Left and right arrows denote runs of ancestral sequence that could be due to a switch-error further away from the starting locus. The haplotypes tagged in blue were removed when selecting per sample the haplotype sharing the most sequence with others.

